# Using Graph Convolutional Neural Networks to Learn a Representation for Glycans

**DOI:** 10.1101/2021.03.01.433491

**Authors:** Rebekka Burkholz, John Quackenbush, Daniel Bojar

## Abstract

As the only nonlinear and most diverse biological sequence, glycans offer substantial challenges for computational biology. These complex carbohydrates participate in nearly all biological processes – from protein folding to the cellular entry of viruses – yet are still not well understood. There are few computational methods to link glycan sequences to functions and those that do exist do not take full advantage of all the available information of glycans. SweetNet is a graph convolutional neural network model that uses graph representation learning to facilitate a computational understanding of glycobiology. SweetNet explicitly incorporates the nonlinear nature of glycans and establishes a framework to map any glycan sequence to a representation. We show that SweetNet outperforms other computational methods in predicting glycan properties on all reported tasks. More importantly, we show that glycan representations, learned by SweetNet, are predictive of organismal phenotypic and environmental properties. Finally, we present a new application for glycan-focused machine learning, the prediction of viral glycan-binding, that can be used to discover new viral receptors and monitor rapidly mutating viruses.

## Introduction

Glycans are complex carbohydrates and are a fundamental biological sequence that is found both as isolated entities as well as covalently bound to proteins, lipids, or other molecules (Varki, 2017). Because of their wide range of interactions, they are intricately involved in protein function (Dekkers et al., 2017; Solá and Griebenow, 2009), cellular function (Parker and Kohler, 2010; Zhao et al., 2008), and organismal function (Haltiwanger and Lowe, 2004; Stanley, 2016). As the only biological sequence with both a non-universal alphabet (consisting of monosaccharides) and nonlinear branching, glycans are highly complex biopolymers. This complexity is further compounded by the fact that glycans are created through a non-templated biosynthesis involving a stochastic interplay of multiple glycosyltransferases and glycosidases (Lairson et al., 2008), such as in the secretory pathway of eukaryotic cells that shapes the range of glycans on secreted proteins (Arigoni-Affolter et al., 2019).

Despite the important role that glycans inhabit in biology, the complexities in their composition and biosynthesis have slowed progress in the experimental and computational study of glycans. Computational approaches to analyzing glycans are mostly limited to counting the occurrence of curated sequence motifs and using this information as input for models predicting glycan properties (Bao et al., 2019; Coff et al., 2020). Recently, deep learning has been applied to the analysis of glycan sequences, creating glycan language models based on recurrent neural networks (Bojar et al., 2020a, 2020b, 2021). The glycan language model SweetTalk views glycans as a sequence of “glycowords” (subsequences that describe structural contexts of a glycan) and was used to predict the taxonomic class of glycans as well as their properties, such as immunogenicity or contribution to pathogenicity. While the usage of glycowords and additional data augmentation strategies in SweetTalk partly accounted for the nonlinear nature of glycan sequences, recurrent neural networks cannot fully capture the branched or tree-like architecture that is seen in most glycans. This implies that alternative model architectures that can fully integrate this nonlinearity should be able to extract more information from glycan sequences, thereby increasing prediction performance.

Advances in deep learning have produced a number of neural network architectures that are capable of analyzing graph- or tree-like structures (Henaff et al., 2015; Wu et al., 2020). These graph neural networks capitalize on the information contained in nodes and their connecting edges as well as the contextual information contained in graph neighborhoods and modules to predict properties of both individual nodes and entire graphs. One of the most useful methods in graph neural networks is message passing by convolutions, a procedure in which a node is described by a combination of the features of surrounding nodes (Henaff et al., 2015; Li and Cheng, 2020). Graph convolutional neural networks (GCNNs) have been used to great effect for studying social networks (Li et al., 2020) or epidemic forecasting (Kapoor et al., 2020) and have also been applied to proteins (Gligorijevic et al., 2019) and small molecule drugs (Nguyen et al., 2020; Torng and Altman, 2019). In the latter, molecules are seen as molecular graphs, with atoms as nodes and bonds as edges. GCNNs also outperform widely used fingerprint-based methods in predicting small molecule properties such as toxicity or solubility (Liu et al., 2019).

SweetNet is a deep learning method that we developed to take advantage of the flexible graph representation structure of GCNNs. SweetNet treats glycan sequences akin to molecular graphs and thereby accounts for their tree-like structure. Viewing monosaccharides and linkages as nodes and their connections as edges allows for the application of GCNNs to glycan sequences without any further manipulations or data augmentation. On a range of reported prediction tasks, we demonstrate that SweetNet yields considerably better prediction results than reported glycan prediction models. We further demonstrate that the latent representations learned by SweetNet are more informative than those derived using other modeling methods. This improved performance is due to the representation of glycans as molecular graphs, a conclusion we also confirm by analyzing structural graph properties of glycans.

We demonstrate the value of SweetNet and the resulting glycan representations in two applications. First, we show that glycans contain information about phenotypic and environmental properties of their organisms that can be extracted via glycan representations. We use this phenomenon to identify phenotypic clusters in the plant order Fabales (dicotyledonous flowering plants that include the legumes), such as having pronounced seeds or fern-like leaves, that are clearly distinguished by their glycans. We further extend this to the kingdom Animalia, identifying clusters of animals inhabiting similar environmental niches (such as amphibians and fish) and revealing the glycan-based placement of *Homo sapiens* apart from most mammals including primates. Our analyses emphasize the important distinction between genomic and glycomic similarity and could enable a new classification of phenotypically or environmentally similar organisms. In particular, this classification may help us to understand how to translate some biological findings from model organisms to the understanding of human systems.

Finally, we show that SweetNet can be used to identify new glycan receptors for viruses by presenting a new glycan-focused prediction task, the prediction of the binding intensities between viral proteins and glycans. For this, we train a SweetNet-based model on a glycan array dataset probing interactions of influenza virus strains and glycans. We demonstrate that this model can then quantitatively predict the glycan-binding behavior of different influenza virus strains. Our model recapitulates known binding specificities of influenza virus and we show that these predictions can be extended to other viruses, such as coronaviruses or rotaviruses. We add to these observations by identifying enriched binding motifs, such as complex motifs from human milk oligosaccharides for rotaviruses, indicating that our model can be used to rapidly identify new glycan receptors for viruses. SweetNet thus represents a new state-of-the-art in glycan-focused machine learning and will enable future investigations into the important roles of glycans.

## Results

### Developing a graph convolutional neural network for glycans

The nonlinear branching structure of glycans, together with their diversity, has hitherto presented an obstacle to the development of machine learning models for glycobiology that fully capitalized on the rich information in glycan sequences. The use of a glycoword-based language model overcame some of these limitations, allowing for the prediction of glycan immunogenicity, pathogenicity, or taxonomic class (Bojar et al., 2020a, 2020b, 2021); data augmentation inspired by graph isomorphism further improved predictions (Bojar et al., 2021). This led us to consider whether the structure of glycans as graphs or trees could be better captured by neural network architectures specifically developed for modeling graphs. Therefore, we developed SweetNet, a graph convolutional neural network (GCNN) that uses the connectivities and identities of monosaccharides and linkages in a glycan as input to predict glycan properties. While linkages might be intuitively interpreted as edges in a graph, we chose to characterize them as nodes. This decision was motivated by the prominence of short glycan motifs, such as the Tn antigen (“GalNAc(α1-“), which otherwise would have been precluded from our analyses.

To find an appropriate model architecture for a GCNN trained on glycan sequences, we chose the task of predicting which species a given glycan sequence came from as the task for building SweetNet. This multiclass classification with 581 unique classes represented one of the most challenging tasks for language models trained on glycan sequences, especially regarding rare classes, and thus offered a suitable challenge for identifying a better model architecture. We constructed neural networks with several different graph convolutional operators, including the simple graph convolutional operator (SGConv) (Wu et al., 2019), the GraphSAGE operator (Hamilton et al., 2018), and k-dimensional graph neural network operators (GraphConv) (Morris et al., 2020). All of the graph convolutional operators we considered outperformed language model-based classifiers, which supported our hypothesis that graph-based models would be more appropriate for branching glycans. Among these, models based on GraphConv operators produced the best models (Table 1).

**Table 1.** Selecting an architecture for a glycan-focused graph convolutional neural network. We trained several machine learning models (random forest, K-nearest neighbor), deep learning models such as the glycan-based language model SweetTalk, and graph convolutional neural networks with different operators (SGConv, SAGEConv, GraphConv) for the prediction of which species a given glycan stemmed from. Mean values from five independent training runs for cross-entropy loss (except for random forest and KNN which do not use this loss function), accuracy, and Matthew’s correlation coefficient on a separate test set are shown. The inclusion of a boom layer and context pretraining is indicated in the model column. For each metric, the superior value is bolded.

| Model | Cross-Entropy Loss | Accuracy | Matthew’s Correlation Coefficient |
| --- | --- | --- | --- |
| Random Forest | - | 0.3025 | 0.2920 |
| KNN | - | 0.3146 | 0.3015 |
| SweetTalk | 3.9550 | 0.3651 | 0.3496 |
| SGConv | 2.5741 | 0.4000 | 0.3882 |
| SAGEConv | 2.5183 | 0.4097 | 0.3989 |
| GraphConv | 2.5173 | 0.4192 | 0.4078 |
| GraphConv<br>Boom | **2.3756** | **0.4430** | **0.4326** |
| GraphConv<br>Boom Pretrain | 2.5048 | 0.4278 | 0.4166 |

We then sought to further enhance model performance by including a boom layer, which has been shown to enhance model performance in other contexts by escaping local minima (Merity, 2019), and observed a further increase in classification performance (Table 1). Hypothesizing that unsupervised pre-training on a larger set of glycans that included glycans without known species labels would further improve performance, we constructed a context prediction task (Hu et al., 2020), in which the model is used to predict the identity of a randomly chosen hidden node, given the connectivities and the other nodes in a glycan. We reasoned that this procedure should allow the model to learn more regularities and context effects from a larger set of diverse glycan sequences. While we were able to successfully pre-train our model, fine-tuning on the species prediction task did not further improve performance, suggesting that this context-dependent information was already incorporated during normal training. Overall, using SweetNet (Figure 1), we achieved an increase of nearly 8% in absolute accuracy for the challenging task of predicting the species of a glycan relative to the previous best method (SweetTalk).

**Figure 1.**
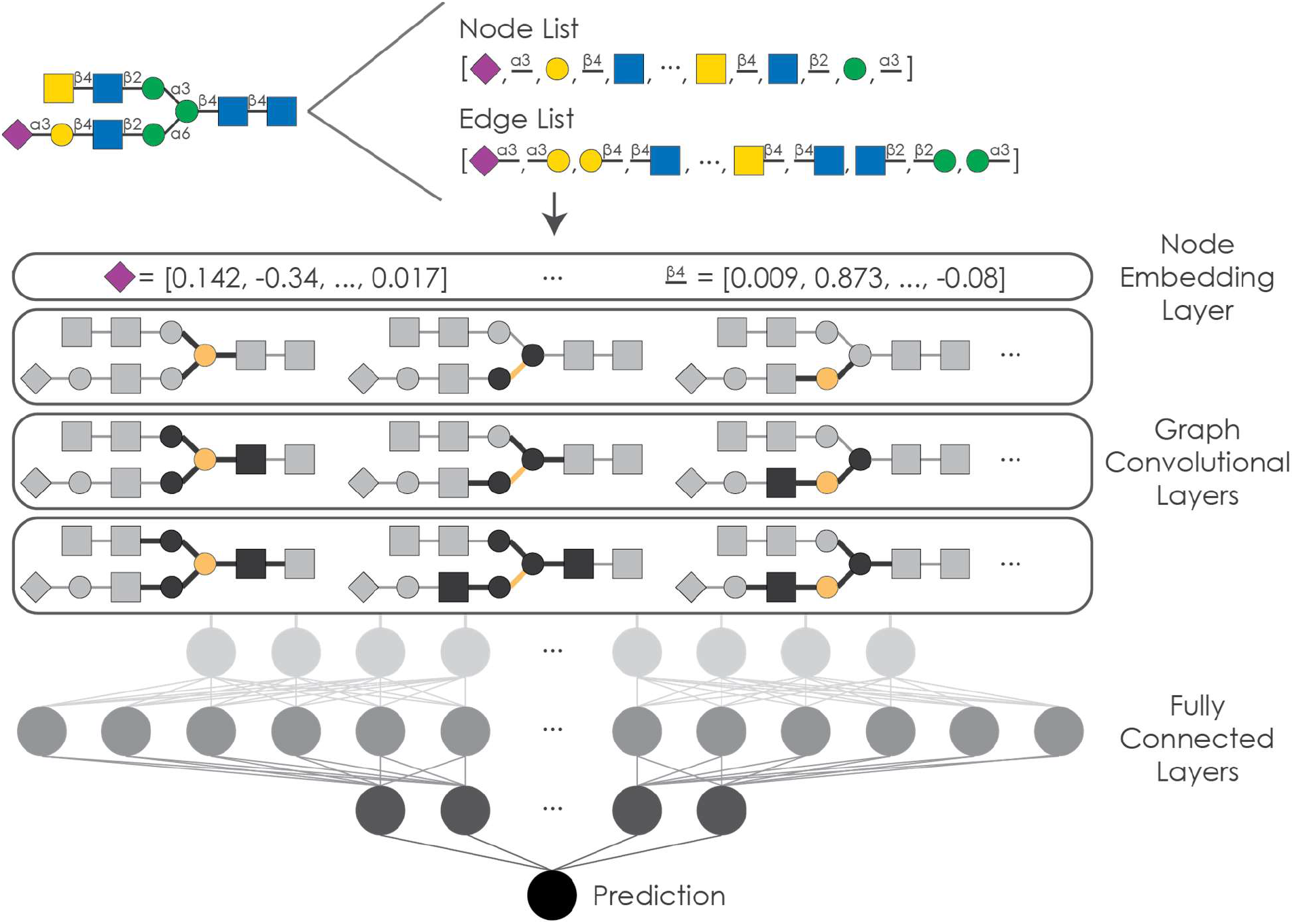
Schematic representation of the graph convolutional neural network SweetNet for analyzing glycans. Glycans are processed into a node list containing all occurring monosaccharides and linkages as well as a list of edge indices detailing the graph connectivity. This information is used as input for SweetNet by generating node features via a representation layer and then feeding the input through three graph convolutional layers. Subsequently, three fully connected layers, including a boom layer, use this information to generate a prediction.

### SweetNet outperforms alternative model architectures on all tasks

Next, we set out to demonstrate that a model architecture focusing on the inherent nature of glycans as molecular graphs is both robust and generalizable. For this, we tested SweetNet on all other prediction tasks that have been previously attempted with glycan-focused machine learning models, predicting higher taxonomic groups of a glycan, predicting glycan immunogenicity, and association with pathogenicity. We found that SweetNet models outperformed all other methods for all prediction tasks (Table 2, Table S1). Compared to previous benchmarks (SweetTalk-based models), SweetNet-based models achieved absolute accuracy increases between 1 and 11 percent (average: 5.16%), depending on the prediction task. This made us confident that GCNNs are a more potent architecture for modeling glycan characteristics and functions.

**Table 2.**
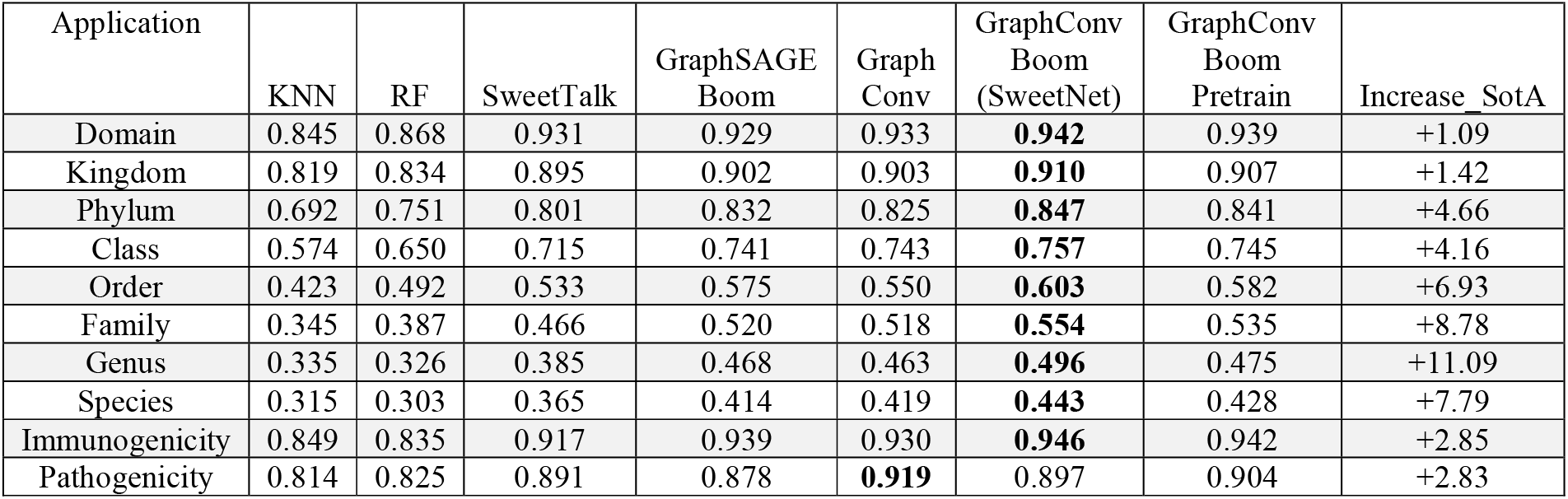
Comparison between different machine learning architectures to extract information from glycans. For a range of prediction tasks (Application), machine learning models (K-nearest neighbor classifier, random forest), deep learning-based language models (SweetTalk), and GCNNs were tested. The choice of graph neural network operator (GraphSAGE, GraphConv) and the presence of a boom layer and pre-training are indicated. Mean accuracy values of five independent training runs for each model on a separate test set are given and Matthew’s correlation coefficient values can be found in Table S1. For each prediction task, the best value is bolded. Performance improvement relative to the previous state-of-the-art model (SweetTalk) is shown in absolute percent increase.

| Application | KNN | RF | SweetTalk | GraphSAGE<br>Boom | Graph<br>Conv | GraphConv<br>Boom<br>(SweetNet) | GraphConv<br>Boom<br>Pretrain | Increase_SotA |
| --- | --- | --- | --- | --- | --- | --- | --- | --- |
| Domain | 0.845 | 0.868 | 0.931 | 0.929 | 0.933 | **0.942** | 0.939 | +1.09 |
| Kingdom | 0.819 | 0.834 | 0.895 | 0.902 | 0.903 | **0.910** | 0.907 | +1.42 |
| Phylum | 0.692 | 0.751 | 0.801 | 0.832 | 0.825 | **0.847** | 0.841 | +4.66 |
| Class | 0.574 | 0.650 | 0.715 | 0.741 | 0.743 | **0.757** | 0.745 | +4.16 |
| Order | 0.423 | 0.492 | 0.533 | 0.575 | 0.550 | **0.603** | 0.582 | +6.93 |
| Family | 0.345 | 0.387 | 0.466 | 0.520 | 0.518 | **0.554** | 0.535 | +8.78 |
| Genus | 0.335 | 0.326 | 0.385 | 0.468 | 0.463 | **0.496** | 0.475 | +11.09 |
| Species | 0.315 | 0.303 | 0.365 | 0.414 | 0.419 | **0.443** | 0.428 | +7.79 |
| Immunogenicity | 0.849 | 0.835 | 0.917 | 0.939 | 0.930 | **0.946** | 0.942 | +2.85 |
| Pathogenicity | 0.814 | 0.825 | 0.891 | 0.878 | **0.919** | 0.897 | 0.904 | +2.83 |

Analogous to the species prediction task, we observed that context pre-training did not improve predictive performance, yet including a boom layer did increase model performance in nearly all cases. Further, even with a boom layer, the performance of GraphSAGE operators was inferior to that of the GraphConv operator. We also observed that, when applied to the task of predicting the taxonomy of a glycan, SweetNet could be trained approximately 30% faster than the equivalent SweetTalk models. SweetNet was also considerably more data-efficient than SweetTalk, surpassing SweetTalk performance even if only a third of the full dataset was used for training (Figure S1). These results confirmed our hypothesis that graph-based models are more efficient in extracting information from glycan sequences than alternative neural network architectures. Further, these data-efficient algorithms could allow prediction even given the relative scarcity of available glycan sequences, caused by the experimental difficulties of working with them.

As we anticipated, challenging prediction tasks with many classes and fewer datapoints per class, such as the prediction of species or genus, saw greater performance gains when using SweetNet than balanced, binary classifications. This is likely the result of a better match between the graph-like nature of glycans and the GCNN architecture, allowing the model to learn more associations and statistical dependencies to use for prediction. Further, the performance of SweetTalk-based models decreased with increased branching while the performance of SweetNet-based models improved with increased branching (Figure S2), indicating that GCNNs such as SweetNet use structural properties unique to graphs, such as the number of branches or various connectivity statistics as features in the classification model.

We next generated a set of graph properties of the glycans in our dataset by calculating various connectivity statistics (Table S2, see Methods for details) and trained a random forest classifier to predict the taxonomic kingdom of a glycan to evaluate whether models could extract information from purely structural features of glycans, without their sequence. Our trained random forest model achieved a predictive accuracy of 61.4% on a separate test set – worse than the random forest model trained on sequence features (Table 2) yet substantially better than random predictions. This confirmed that graph properties by themselves are informative for predicting glycan properties, albeit possibly less so than sequence features.

Analyzing the feature importance of this model, we observed that the most important graph feature for a kingdom-level classification was the number of node types in a glycan (reflecting overall glycan sequence diversity; Figures S3A-B), which we also observed to be important for predicting the contribution of glycans to pathogenicity (Figure S3C), the immunogenicity of a glycan (Figure S3D), and the type of glycan (Figure S3E). To our surprise, we found that a smaller number of node types (meaning greater homogeneity in the glycan) predicted higher immunogenicity (Figures S3F-G); this result could be due to the presence of multiple binding sites for antibodies and the innate immune system. Other important features for predicting glycan immunogenicity and class for instance included aspects of the harmonic centrality, which is related to a graph’s compactness (Figures S3C-E).

### Glycan representations learned by SweetNet are more informative compared to alternative models

An additional advantage of deep learning models is that, during training, representations of glycans are learned that can then be used for visualization and downstream prediction tasks. Reasoning that a model with superior prediction performance should have also learned more informative representations, we extracted glycan representations for some of the SweetNet models we trained. For this, we used glycan sequences as input for the trained model and extracted the results from the graph convolutional layers, directly prior to the fully connected part of the network. These representations can be visualized in two dimensions to identify clusters in the data. We demonstrated this with the example of the SweetNet-based model predicting glycan immunogenicity, that resolved clear clusters of N- and O-linked glycans as well as glycolipids, respectively (Figure 2A). Further, the relative proximity of O-linked glycans and glycolipids to immunogenic bacterial glycans is consistent with phenomena of molecular mimicry (Bojar et al., 2021; Carlin et al., 2009), indicating that SweetNet incorporates biologically meaningful information.

**Figure 2.**
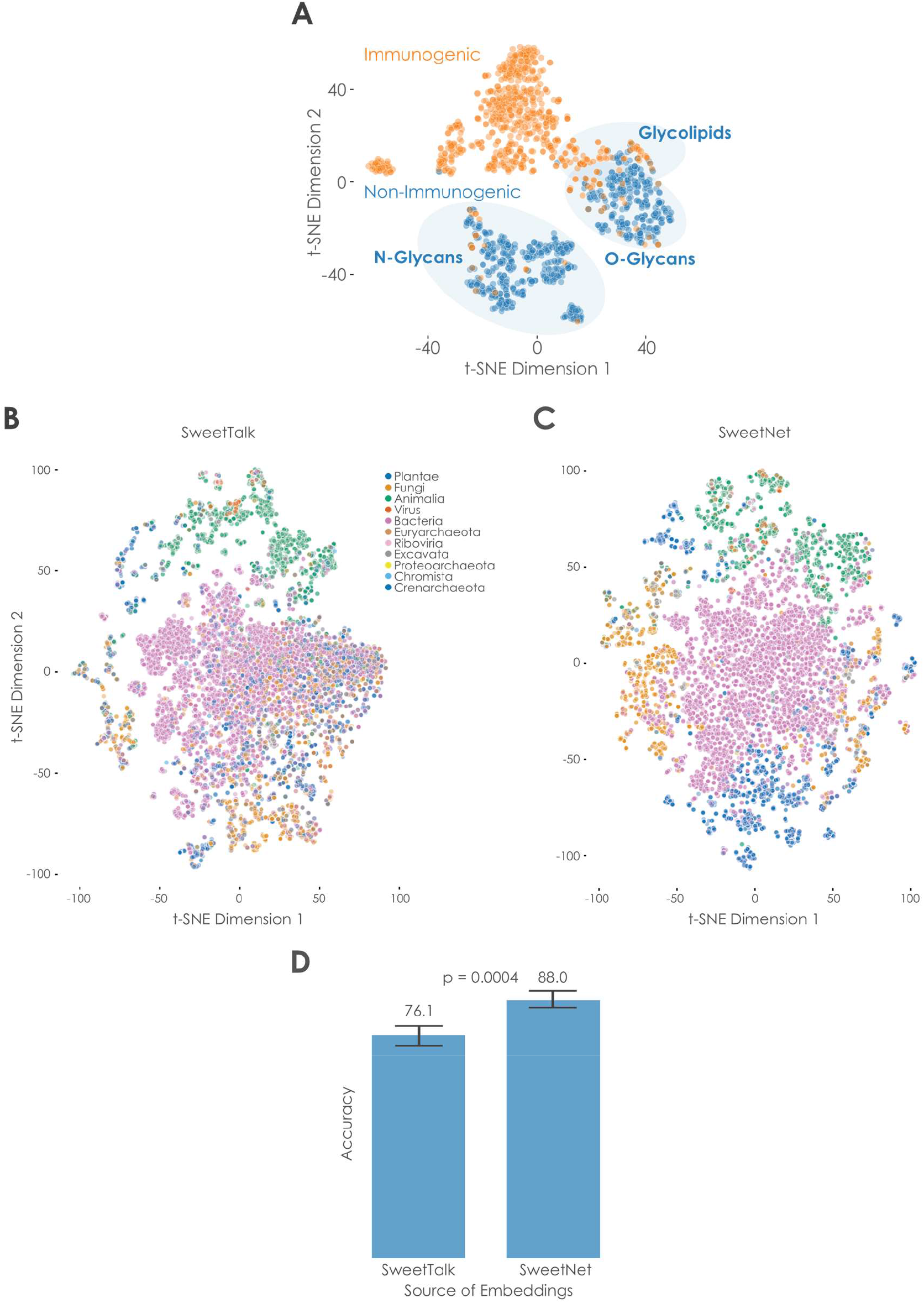
Comparison of glycan representations obtained by machine learning. **(A)** Immunogenic glycan representations learned by SweetNet. Glycan representations for all glycans with immunogenic information were extracted from a trained SweetNet-based model and are shown via t-SNE (van der Maaten and Hinton, 2008), colored by their immunogenicity label and annotated by glycan classes. (**B-C)** Taxonomic glycan representations learned by the SweetTalk and SweetNet. Glycan representations for all glycans with taxonomic information in our dataset were generated by SweetTalk (**B**) and SweetNet (**C**) trained on predicting the taxonomic genus a given glycan stemmed from. These representations are shown via t-SNE and colored by their taxonomic kingdom. (**D)** Comparing information value of representations obtained by SweetTalk and SweetNet. Logistic regression models were trained on the representations obtained from the genus-level SweetTalk and SweetNet models in order to predict the taxonomic kingdom of a glycan. The achieved accuracy from representations from five training runs, respectively, is shown here and was compared between models by a Welch’s t-test.

As we observed the greatest performance increase of SweetNet relative to SweetTalk models in the taxonomic task of genus prediction, we compared glycan representations from SweetNet and SweetTalk models trained on this task (Bojar et al., 2021) (Figures 2B-C). Coloring glycans by kingdom allowed us to observe that SweetNet-based representations improved separation of glycans into taxonomic kingdoms, with clearer neighborhood separations; this can also be seen in the higher adjusted Rand index when K-means clustering kingdoms from SweetNet-based representations (0.203) than from SweetTalk-based representations (0.146).

To further quantify the information in these representations, we trained logistic regression models to predict the taxonomic kingdom of a glycan from its representation gained by genus-level SweetTalk or SweetNet, respectively. The SweetNet representations again demonstrated superior performance (Figure 2D; ~88% accuracy) compared to the model trained on SweetTalk representations (~76% accuracy), indicating the value of SweetNet representations for downstream analyses. The accuracy achieved by using representations from the genus-level SweetNet model even reached the level obtained by the kingdom-level SweetTalk model (Table 1), suggesting that cross-training for hierarchically related tasks could provide additional predictive power (Sarawagi et al., 2003). Additionally, representations learned by SweetNet recovered clusters of glycans with certain graph properties, such as high number of node types or high average eigenvalues (Figure S4).

### Extracting Phenotypic and Environmental Properties from Glycan Representations

Considering the rich glycan representations learned by SweetNet, we set out to improve on reported glycan-based phylogenies (Bojar et al., 2020a). Conveying phenotypic plasticity and covering every cellular surface, glycans are a major driver of evolution (Lauc et al., 2014) and mediate organismal properties more directly than DNA. Thus, a glycan-based phylogeny could offer new insights into evolutionary histories and phenotypic similarity between species beyond that seen through DNA analysis. We constructed a proof-of-principle dendrogram of all species in the order Fabales with known glycans by averaging their glycan representations, constructing a cosine distance matrix, and performing hierarchical clustering. This revealed a clear clustering of taxonomic groups, with close association of species in the same genera and families, enabling us to establish a glycan-based phylogeny (Figure 3A).

**Figure 3.**
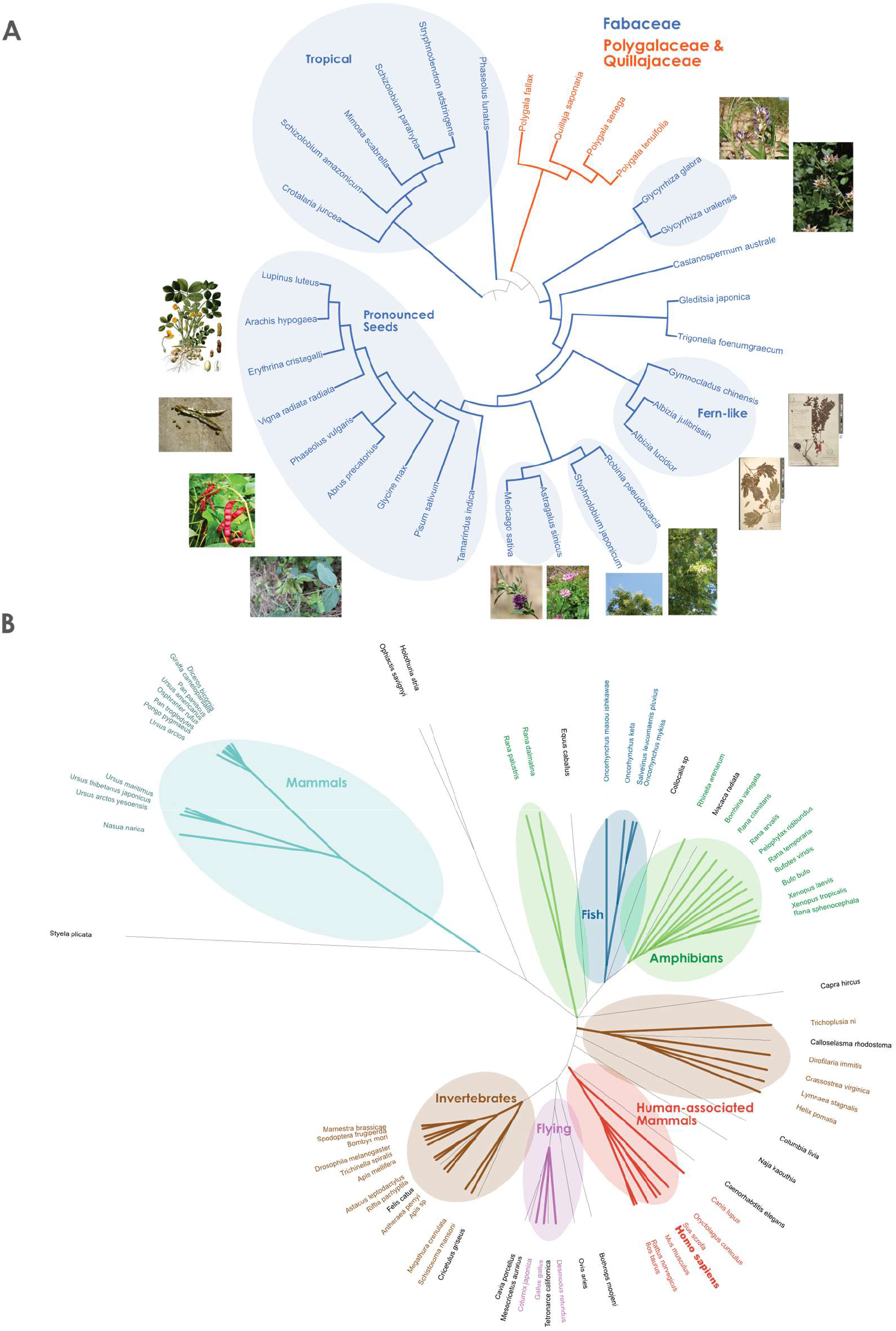
Glycan-based phylogenetic trees. For all 30 species from the order Fabales with at least five glycans (**A**) or 93 species from the kingdom Animalia with at least two glycans in our dataset (**B**), we averaged their glycan representations from the species-level SweetNet model, constructed a cosine distance matrix between all species, and performed hierarchical clustering to obtain a dendrogram. The shown phylogenetic tree was drawn with the Interactive Tree of Life v5.5 software (Letunic and Bork, 2019). For **A**, we colored species belonging to the taxonomic families Fabaceae and Polygalaceae / Quillajaceae, respectively. We further annotated clusters enriched for certain groups of plants (**A**) or animals (**B**) that shared characteristics. The species *Homo sapiens* is depicted in bold.

In the dendrogram, we found clusters of plant species that share environmental and/or phenotypic similarities. This includes a cluster of plants occurring in tropical environments (South America and Africa), a cluster containing Fabaceae species that produce pronounced, mostly edible, seeds such as *Arachis hypogaea* (peanut) or *Glycine max* (soybean), and smaller clusters characterized by plants with similar leaf or flower phenotypes (Figure 3A). These results indicate that glycans carry considerable information about phenotypic and environmental characteristics of species, and can be used to group species with shared properties and establish the notion of glycan-based relatedness, including the known effects of glycan-mediated phenotypic plasticity (Lauc et al., 2014).

We applied this method to all available species of the kingdom Animalia (Figure 3B) to draw a glycan-based tree of (animal) life. While differences in coverage prevented us from including all known species and distorted some relationships, we still were encouraged to see meaningful patterns emerge from this glycan-based phylogeny. We observed adjacent clusters for amphibians and fish, potentially reflecting their overlapping environmental range or their evolutionary history. We also find distinct clusters for flying animals (birds and bats), invertebrates, and mammals, respectively. It is interesting that *Homo sapiens* was absent from the cluster that included primates and instead clustered with, perhaps tellingly, animals that surround us, such as pigs, cows, mice, rats, rabbits, and dogs. The most closely associated species in the cluster, the pig *Sus scrofa,* is a prime candidate for xenotransplantation and is a major source of tissue for heart valve transplants (Burlak et al., 2013; Manji et al., 2015). These clustering results suggest that environmental factors influence glycan production and structure in ways that are distinct from what we might expect based on DNA phylogeny. Some of these glycan alterations may reflect specific genomic changes as there are reports about several gene losses in the evolutionary history of *Homo sapiens* that have changed the glycome of our species (Irie et al., 1998; Lanteri et al., 2002).

### Using SweetNet to explain virus-glycan binding

Most viruses bind to glycan receptors before, in some cases, transitioning to proteinaceous receptors during cell entry (Koehler et al., 2020; Thompson et al., 2019). The specificity and affinity of these glycan binding events are essential for both virulence and host specificity, such as for influenza virus strains which usually prefer either α2-3- or α2-6-linked sialic acids in their glycan receptors (Viswanathan et al., 2010). We constructed a model that, given a viral protein sequence and a glycan, could predict their interaction in hopes of better understanding viral cell entry, developing methods to monitor emerging viral strains, and to suggest glycan-based antivirals.

Our model comprised a recurrent neural network analyzing the protein sequence, a module analyzing the glycan sequence (see below), and a fully connected part concatenating the results of both prior modules to predict the binding intensity of a protein-glycan pair. Using influenza virus and its glycan-binding protein hemagglutinin as a test case, we gathered 126,894 measured interactions between hemagglutinin variants with available sequences and glycans from the glycan array database of the Consortium for Functional Glycomics (Gao et al., 2019) (Table S3). Next, we determined which module for analyzing glycan sequences led to the highest prediction performance. For this, we tested three different modules: (1) a fully connected neural network that used the counts of mono-, di-, and trisaccharides of glycan sequences as input, (2) a SweetTalk-based glycan language model, and (3) a SweetNet-based GCNN that we introduced here. The SweetNet-based approach again yielded an improved performance over the currently available sequence motif-based and language model-based approaches (Table 3, Table S1).

**Table 3.**
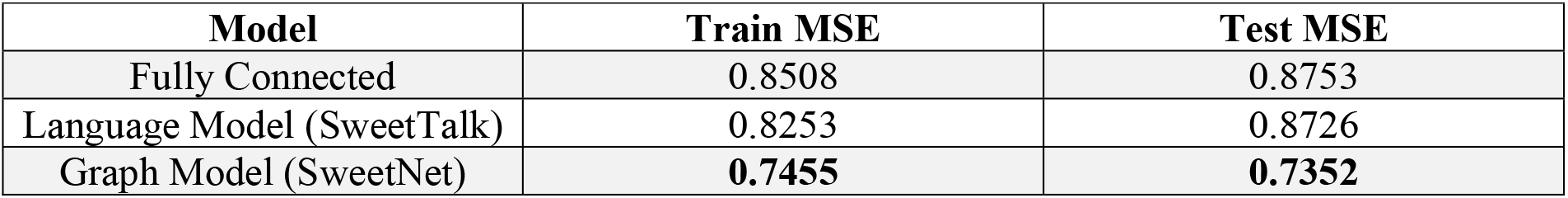
Developing a model predicting viral glycan-binding behavior. Models consisted of a recurrent neural network analyzing the protein sequences of viral hemagglutinin as well as either a fully connected neural network using the counts of mono-, di-, and trisaccharides as input (Fully Connected), a SweetTalk-based glycan language model, or a SweetNet-based GCNN. All models were trained to predict Z-score transformed glycan-binding of hemagglutinin from various influenza virus strains. Average mean square errors (MSE) from five independent training runs, from both the training and independent test set, are shown here. Superior values for each metric are bolded.

We then went on to ensure that this mean performance implied that most predictions would fall within this level of prediction error. For this, we collected all residuals between observed and predicted Z-scores (Figure 4A), indeed observing that most residuals fall within the margin described by the mean prediction error. Further, the performance of our model remained stable for subgroups, such as virus subtypes or host organisms (Figure 4B).

**Figure 4.**
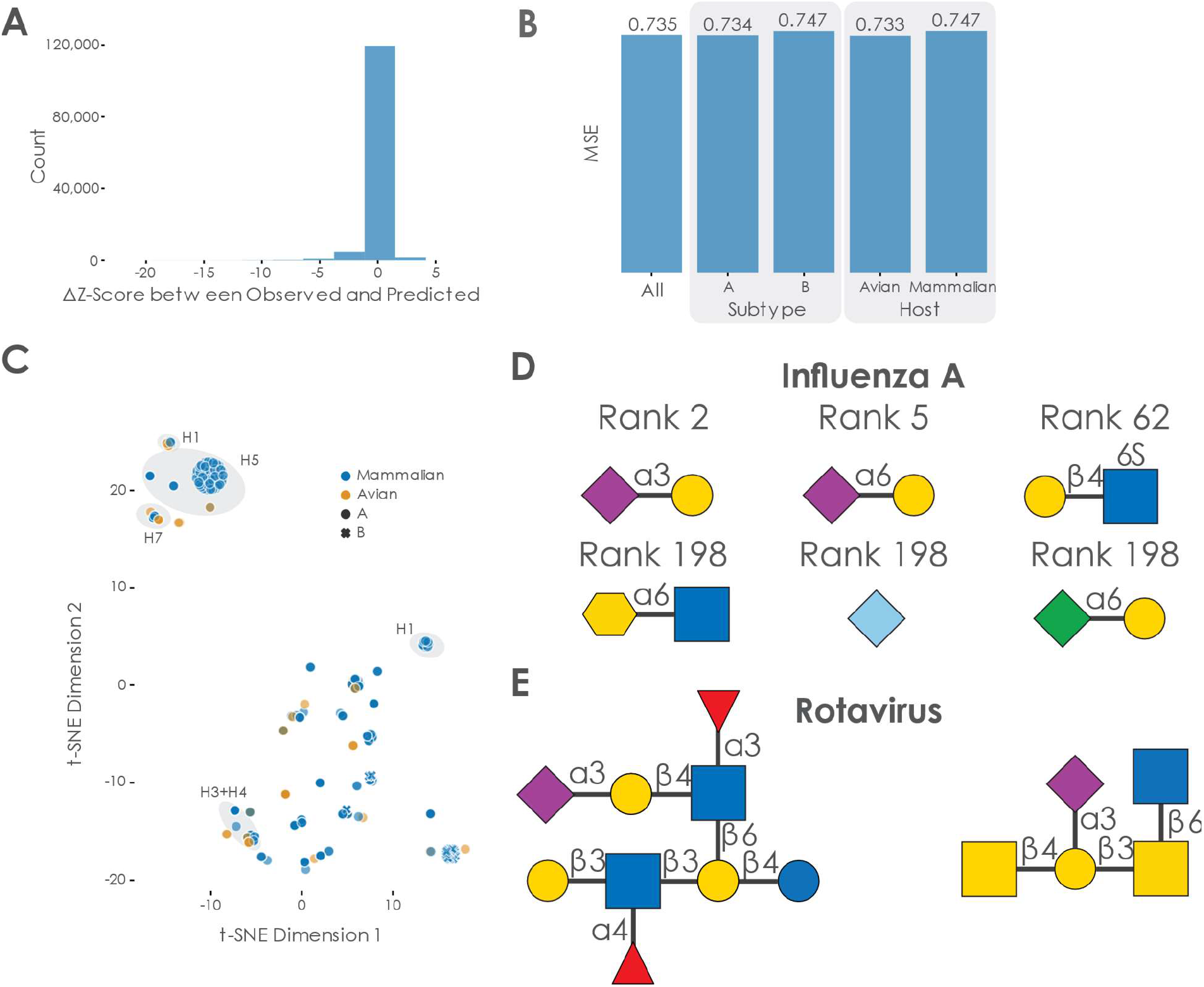
Characterizing the SweetNet-based model predicting virus-glycan binding. **(A)** Distribution of residuals between predicted and observed Z-scores. We calculated the difference between all observed and predicted Z-scores of hemagglutinin-glycan binding interactions which is shown here as a histogram. (**B)** Analyzing model performance on subgroups in the data. We obtained the mean square error (MSE) of our trained model for subgroups such as virus subtype or host organism in our data. (**C)** Hemagglutinin representations learned by the protein-analyzing module of the model. We obtained the representation learned by our protein-analysis module from the last state of our LSTM for all 339 unique protein sequences in our dataset. These representations are shown via t-SNE and colored / marked by their host organisms and virus subtype, respectively. Clusters of hemagglutinin subtypes are further annotated. (**D)** Examples of glycan motifs relevant for the prediction of hemagglutinin-glycan binding by SweetNet-based models. Some of the glycan motifs that were significantly relevant for prediction are shown in the SNFG nomenclature, together with their median rank across host species. (**E)** Examples of glycans with high predicted binding to rotavirus.

Next, we analyzed the representations learned by the protein-analyzing LSTM module in our model, observing clustering according to the hemagglutinin subtype (Figure 4C). Additionally, while we did observe the separation of hemagglutinin from influenza A and B, the split between mammalian and avian hemagglutinin was less obvious, even though the classical view is that they differ considerably in their glycan-binding specificity. Supporting our findings, systematic binding studies have indeed identified several avian influenza hemagglutinin subtypes, such as H4 and H9, that exhibit binding properties similar to mammalian influenza hemagglutinin (Shelton et al., 2011).

We then identified glycan motifs that are important for the binding of hemagglutinin. Most approaches apply some form of subtree frequency mining to the glycan array data (Coff et al., 2020), identifying preferentially bound glycan fragments (Cholleti et al., 2012). However, we wanted to capitalize on the predictive nature of our trained model and used all our 19,775 available glycan sequences, which are to the most part not covered by existing glycan arrays, as inputs to the trained model for each host species. Then, we analyzed the resulting binding predictions to ascertain which glycan motifs were most predictive for high-affinity binding across species (Figure 4D, see Methods for further details). Assaying all 23,170 observed glycan motifs with lengths 1, 2, or 3 confirmed results achieved with standard methods, with Neu5Ac as the most important motif (median rank 1), and several known binding motifs such as Neu5Ac(α2-3)Gal (rank 2) or Neu5Ac(α2-6)Gal (rank 5; see Table S4 for all motifs). Published studies also suggest that sulfated glycan motifs may serve as binding motifs for influenza hemagglutinin (Ichimiya et al., 2014), which is consistent with our results (significant motifs with median ranks between 52 and 198; see Table S4).

We observed that the SweetNet-based model had learned relevant chemical information to predict hemagglutinin-glycan interactions by focusing on negatively charged motifs (containing carboxylates, sulfates, or phosphates) and sialic acids and structurally related monosaccharides in particular (Neu5Ac, Neu5Gc, Kdn, Kdo). This is especially encouraging as monosaccharides such as Kdo (3-Deoxy-d-manno-oct-2-ulosonic acid), a bacterial analogue of Kdn, were not present in the dataset used to train the model. This indicates that the model learns general features of glycans that are predictive of properties of “novel” glycan motifs. Although Kdo has, to our knowledge, not yet been proposed to bind influenza hemagglutinin, SweetNet suggests that it could be a target for influenza hemagglutinin.

We next wanted to test the generalizability of SweetNet for predicting other viral targets. For this, we trained a SweetNet model on a dataset combining the influenza virus glycan arrays with data from 83 additional glycan arrays that had been probed with a wide range of viruses (validation MSE: 0.784; Table S5). We then predicted the most important binding motifs for coronaviruses with this model and identified sialic acid motifs as well as sulfated glycan motifs (Table S6). These results are supported by reported binding motifs (Milewska et al., 2014) that naturally occur in glycosaminoglycans such as heparan sulfate. Next, we used the model to identify glycans with high predicted binding to rotaviruses, a common neonatal virus. Among the top 10 predicted glycans, we observed, next to expected sialic acid motifs, core 2 O-glycans (Figure 4E) that were indicated to bind rotaviruses (Pang et al., 2018). We additionally identified a glycan containing the Gal(β1-3)GlcNAc(β1-3)Gal(β1-4)Glc motif (Figure 4E, Table S7) that was previously described to be a potential decoy receptor from human milk that is bound by rotaviruses (Yu et al., 2014). Crucially, this glycan was not part of our training dataset and the authors of this previous study had to use a specialized array made from human milk oligosaccharides to discover this binding motif. This demonstrates that SweetNet-based models can make useful predictions for unobserved glycans and positions our virus-glycan binding model to rapidly make predictions of bound glycan receptors for emerging virus variants.

## Discussion

We tend to think of nucleic acids as the most important biological molecules because they store the genetic code and facilitate protein synthesis, or proteins themselves because of their roles in mediating cellular processes and functions and serving in structural roles. However, cells are supported by a vast network of interacting biological molecules that also include lipids and complex carbohydrates that serve in essential functional, metabolic, or structural capacities. Glycans represent a unique class of biological molecules in that they have nonlinear, branching structures that allow them to carry out a wide range of functions, encompassing protein folding and degradation, stress response, cell-cell interactions, cell migration patterns, self/non-self discrimination, and microbiome development, composition, and health (Varki, 2017).

Not surprisingly, glycans differ between species and change in response to environmental perturbations and so have the potential to allow us to understand genetic and environmental interactions (Lauc et al., 2014; Springer and Gagneux, 2016). Ideally, one would like to use glycan sequences to gain insights into phenotypic and environmental properties and to predict processes mediated by glycans such as viral infection. However, such applications are still rare, which may be due to the complex structure of glycans and the importance of these structures in determining glycan function.

Graph convolutional neural networks (GCNNs) are a machine learning method that performs convolutional methods on the input graph itself (Henaff et al., 2015; Wu et al., 2020), with structure and features unchanged, rather than creating a lower-dimensional representation. Since the graph retains its original form, the relational inductive bias that is possible is much stronger. Given that glycans can be represented as complex graphs, we believed that GCNNs represented an ideal tool for applications involving glycan-based classification.

SweetNet is a GCNN implementation that fully leverages the tree-like structure of branched glycans. SweetNet-based models of glycans can be trained faster and are considerably more data-efficient than models using other neural network architectures and SweetNet outperforms these models in all of the tasks we analyzed. The data efficiency of SweetNet is an important metric because generating glycan data is not yet as easy or high-throughput as either DNA sequencing or proteomics, meaning this method can take advantage of the relatively sparse existing data available now and that its performance will improve as more data become available.

In using glycan profiles from a wide range of species, we demonstrated that SweetNet can find taxonomic clusters that appear to group species based on both their evolutionary relationships and the ecological niche they inhabit, a finding consistent with the known genetic and environmental effects on glycan synthesis (Springer and Gagneux, 2016). Analyzed in this way, glycans could provide a unique window into similarities between species and their environments and may shed light on the role of glycans in phenotypic plasticity and evolution (Lauc et al., 2014).

By choosing to consider both monosaccharides and linkages as nodes in our glycan graphs, we avoided a limitation of GCNNs that only glycans with a minimum length of at least two monosaccharides (disaccharides or larger) could be analyzed. Thus, SweetNet-based models can analyze important glycan structures such as the Tn antigen (“GalNAc(α1-“), which were inaccessible with previous language model-based approaches, extending the potential applications for glycan-focused machine learning. Additionally, for applications involving these short glycans, the respective node representation learned by SweetNet could be used to extract information.

Ambiguities in nomenclature have spawned a plethora of different formats for describing glycans, yet most formats are either not human-readable or insufficiently convey the branched structure of glycans. Depicting glycans as graphs circumvents these difficulties and offers the most promising nomenclature for predictive models in glycobiology. It also readily facilitates glycoengineering efforts (Kightlinger et al., 2020) by adding or removing nodes and the corresponding edges to a glycan graph and querying trained models for the predicted properties of the proposed glycans. Promising glycans could then be prioritized as design targets for antivirals or other purposes. Further, existing analysis modalities of glycan substructures or subsequences (Bao et al., 2019) could readily be applied to the subset of glycans with high prediction scores to rationalize model predictions. Our trained models predicting virus-glycan binding could also be used to obtain relevant glycan binding motifs to design glycan-based antivirals and additionally assess the potential of designed antiviral candidates by predicting their binding. As this represents only one of the areas of application for our platform, we envision a place in the design-build-test cycles of glycoengineering efforts for our SweetNet-based models.

Glycans have hitherto been neglected in most biological phenomena, at least in part because of the difficulties to work with and analyze glycans. The increasing number of applications in which glycan-focused machine learning has been shown to be feasible bodes well for finally lifting this analysis bottleneck and incorporate glycans into common analysis workflows. This is particularly emphasized by the development of new model architectures and analysis platforms that are more data-efficient, broadening the range of possible applications. Here, we advance both aspects, contributing new applications for glycan-focused machine learning, with our virus-glycan binding predictions, and new, data-efficient models with our GCNN, SweetNet, that can already achieve state-of-the-art performance with small datasets. Our workflows are robust as well as rapid and we envision the application of SweetNet-based models to many glycan-focused classification tasks.

## Contributions

D.B. conceived the method. D.B. and R.B. designed and performed the experiments. D.B., J.Q., and R.B. wrote and edited the manuscript.

## Methods

### Resource availability

#### Lead contact

Communication should be directed to the lead contact, Daniel Bojar

### Materials availability

This study did not generate new unique reagents.

### Data and code availability

Data used for all analyses can be found in the supplementary tables. All code and trained models can be found at https://github.com/BojarLab/SweetNet

### Method details

#### Data processing

For comparing SweetNet to previously reported models, the data used in this study were largely from previous work (Bojar et al., 2021) and consisted of glycan sequences with their associated labels, such as taxonomic class, immunogenicity, or pathogenicity. For the model predicting viral glycan binding, we additionally obtained data from 587 glycan array screens from the Consortium for Functional Glycomics that measured the glycan binding behavior of hemagglutinin from various strains of influenza virus. For each array, we transformed the data to Z-scores. All data can be found in Table S3. For our expanded dataset, we also included Z score-transformed data from 83 arrays testing various viruses (Table S5).

For traditional machine learning models, we generated count variables that detailed the frequency of observed mono-, di-, and trisaccharide motifs in glycans and used these as input features. For glycan-based language models, we followed the data processing detailed previously (Bojar et al., 2021). Briefly, we extracted “glycowords” (trisaccharides in the IUPAC condensed bracket notation) from isomorphic variants of the bracket notation of a glycan and used these as input for a bidirectional recurrent neural network. For graph convolutional neural networks, we converted glycans from the bracket notation to graphs by generating a node list, in which every monosaccharide or linkage constituted a node, and a list of edge indices that detailed the graph connectivity.

For the model predicting viral glycan-binding, we selected a hemagglutinin core sequence that incorporated relevant binding loops (amino acid position 50 to 300) when the full sequence was available, to facilitate comparison to screens in which only partial sequences were available. Then, we label-encoded single amino acids in these sequences and used them as input for a recurrent neural network. This information was then combined with analyses of the corresponding glycan sequences by either 1) motif counting for a fully connected neural network, 2) a language model based on a recurrent neural network, or 3) a graph convolutional neural network.

#### Model training

All models were trained with PyTorch (Paszke et al., 2019) on a single NVIDIA® Tesla® K80 GPU and the architecture as well as hyperparameters were optimized by minimizing the respective loss function via cross-validation of the training set. For all model applications, we used a random split into 80% training and 20% test data. As in previous work (Bojar et al., 2021), we used a stratified split for the taxonomic classifiers to ensure that all classes are split according to this ratio and also only consider classes with at least five known glycans. For the language models, all glycans were brought to the same length by padding. Language models were initialized using Xavier initialization (Glorot and Bengio, 2010) and GCNNs were initialized with a sparse initialization using a sparsity of 10%.

The final SweetNet model consisted of a 128-dimensional node representation layer followed by three iterations of graph convolutional layers, leaky ReLUs, Top-K pooling layers, and both global mean and global maximum pooling operations. The results from these three iterations were added and passed to a set of three fully connected layers interspersed with batch normalization layers, dropout layers, and leaky ReLUs as activation functions. For the final layer, we used a multisample dropout scheme (Inoue, 2019).

All models used a batch size of 32 for training and testing. The ADAM optimizer was used in all cases with a weight decay value of 0.001, together with a starting learning rate of 0.0005 that was decayed according to a cosine function over 80 epochs. Training proceeded for 100 epochs and was stopped early if the loss function did not decrease for at least 20 epochs. Depending on the application, we used binary cross-entropy, cross-entropy, or mean squared error loss functions.

#### Assessing the predictive value of structural graph features in glycans

We extracted 42 different graph features from each glycan to assess the predictive value of the graph representation from a structural point of view (Table S2). These features comprised the number of nodes, the number of different node types, the diameter of each graph (i.e., the maximum shortest path), its branching number (i.e., the number of nodes with 3 or more neighbors), the number of leaves (i.e., number of nodes with only one neighbor), and statistics on different centrality measures. Node and edge centrality measures assign a value to each node or edge in a graph to measure their respective importance. To define the same number of features for each graph with varying numbers of nodes, we always extracted the maximum value, the minimum value, the mean, and the variance across all nodes or edges. All considered centralities are implemented in the Python package NetworkX 2.5 (Hagberg et al., 2008). Specifically, we included degree centrality, betweenness centrality, flow centrality for nodes and edges, eigen centrality, closeness centrality, harmonic centrality, second order centrality, and load centrality. Usually, the degree (i.e., the number of neighbors a node has) plays a major role when comparing different graphs. As it also measures different aspects of branching, we included additional related features in our analysis: the degree assortativity, the number of nodes with at least four neighbors, and the maximum and mean number of leaves a node is connected to, which describes branching at potential binding sites. Furthermore, we added the maximum size of a k-core (comparing different values of k) and a corona to our set of features. These evaluate whether highly connected nodes tend to clique together. The most noteworthy features are the harmonic, flow, and second order centralities, which identify nodes as central that are close to most other nodes in the graph and are visited consistently by random walkers. Intuitively, our derived features are thus related to how compactly a graph is organized.

#### Identifying glycan motifs predictive for hemagglutinin-binding

All glycans in Table S8 were used as input for the trained SweetNet-based model predicting hemagglutinin-glycan binding. This was performed separately for hemagglutinin sequences from viruses of all host species (human, pig, dog, horse, bat, seal, duck, chicken, turkey, shorebird, gull). We then extracted all 23,170 glycan motifs of lengths 1, 2, and 3 from the glycan sequences that were observed in our dataset, as reported previously (Bojar et al., 2021). For each species, we then defined “predicted binding” as a predicted Z-score above 1.645, as suggested by related work on glycan array data (Cholleti et al., 2012). For each glycan motif, we performed a one-tailed Welch’s t-test to ascertain whether glycans containing this motif were more prevalent among glycans predicted to bind. The resulting p-values were subsequently corrected for multiple testing by a Holm-Šidák correction. To gauge the overall relevance of the significant glycan motifs, we then calculated the median rank of each significant motif across all host species.

#### Quantification and statistical analysis

For statistical analysis, this study used Welch’s t-tests with a Holm-Šidák correction for multiple testing correction. All experimental details can be found in the Methods section.

#### Images

Most images used for annotating the glycan-based phylogenetic tree of Fabales (Figure 3A) stemmed from the public domain. Exceptions were from Creative Commons licenses that required attribution and included *Astragalus sinicus* (https://commons.wikimedia.org/wiki/File:Chinese_milkvetch_Ziyunying.JPG), *Glycyrrhiza uralensis* (https://commons.wikimedia.org/wiki/File:Glycyrrhiza_uralensis_IMG_1086.jpg), and *Glycyrrhiza glabra* (https://commons.wikimedia.org/wiki/File:Glycyrrhiza_glabra_Y13.jpg).

### Supplementary Tables

**Supplementary Table 1. Model performances on glycan-focused tasks.** Related to Table 2.

**Supplementary Table 2. Graph features of glycans in dataset.** Related to Figure 2.

**Supplementary Table 3. Glycan array data for influenza virus hemagglutinin.** Related to Figure 4.

**Supplementary Table 4. Enriched glycan motifs for binding influenza virus hemagglutinin.** Related to Figure 4D.

**Supplementary Table 5. Glycan array data for various viruses.** Related to Figure 4.

**Supplementary Table 6. Enriched glycan motifs for binding coronavirus.** Related to Figure 4.

**Supplementary Table 7. Binding predictions for glycans binding to rotavirus.** Related to Figure 4E.

**Supplementary Table 8. All available glycan sequences for identifying enriched motifs.** Related to STAR Methods.

## References

Arigoni-Affolter, I., Scibona, E., Lin, C.-W., Brühlmann, D., Souquet, J., Broly, H., and Aebi, M. (2019). Mechanistic reconstruction of glycoprotein secretion through monitoring of intracellular N-glycan processing. Sci. Adv. 5, eaax8930.

Bao, B., Kellman, B.P., Chiang, A.W.T., York, A.K., Mohammad, M.A., Haymond, M.W., Bode, L., and Lewis, N.E. (2019). Correcting for sparsity and non-independence in glycomic data through a systems biology framework. bioRxiv doi: 10.1101/693507.

Bojar, D., Powers, R.K., Camacho, D.M., and Collins, J.J. (2020a). SweetOrigins: Extracting Evolutionary Information from Glycans. bioRxiv doi: 10.1101/2020.04.08.031948.

Bojar, D., Camacho, D.M., and Collins, J.J. (2020b). Using Natural Language Processing to Learn the Grammar of Glycans. bioRxiv doi:10.1101/2020.01.10.902114.

Bojar, D., Powers, R.K., Camacho, D.M., and Collins, J.J. (2021). Deep-Learning Resources for Studying Glycan-Mediated Host-Microbe Interactions. Cell Host & Microbe 29, 132–144.e3.

Burlak, C., Bern, M., Brito, A.E., Isailovic, D., Wang, Z.-Y., Estrada, J.L., Li, P., and Tector, A.J. (2013). *N* - linked glycan profiling of GGTA1/CMAH knockout pigs identifies new potential carbohydrate xenoantigens. Xenotransplantation 20, 277–291.

Carlin, A.F., Uchiyama, S., Chang, Y.-C., Lewis, A.L., Nizet, V., and Varki, A. (2009). Molecular mimicry of host sialylated glycans allows a bacterial pathogen to engage neutrophil Siglec-9 and dampen the innate immune response. Blood 113, 3333–3336.

Cholleti, S.R., Agravat, S., Morris, T., Saltz, J.H., Song, X., Cummings, R.D., and Smith, D.F. (2012). Automated Motif Discovery from Glycan Array Data. OMICS: A Journal of Integrative Biology 16, 497–512.

Coff, L., Chan, J., Ramsland, P.A., and Guy, A.J. (2020). Identifying glycan motifs using a novel subtree mining approach. BMC Bioinformatics 21, 42.

Dekkers, G., Treffers, L., Plomp, R., Bentlage, A.E.H., de Boer, M., Koeleman, C.A.M., Lissenberg-Thunnissen, S.N., Visser, R., Brouwer, M., Mok, J.Y., et al. (2017). Decoding the Human Immunoglobulin G-Glycan Repertoire Reveals a Spectrum of Fc-Receptor-and Complement-Mediated-Effector Activities. Front. Immunol. 8, 877.

Gao, C., Wei, M., McKitrick, T.R., McQuillan, A.M., Heimburg-Molinaro, J., and Cummings, R.D. (2019). Glycan Microarrays as Chemical Tools for Identifying Glycan Recognition by Immune Proteins. Front. Chem. 7, 833.

Gligorijevic, V., Renfrew, P.D., Kosciolek, T., Leman, J.K., Cho, K., Vatanen, T., Berenberg, D., Taylor, B., Fisk, I.M., Xavier, R.J., et al. (2019). Structure-Based Function Prediction using Graph Convolutional Networks. bioRxiv doi:10.1101/786236.

Glorot, X., and Bengio, Y. (2010). Understanding the difficulty of training deep feedforward neural networks. In Proceedings of the Thirteenth International Conference on Artificial Intelligence and Statistics, pp. 249–256.

Hagberg, A.A., Schult, D.A., and Swart, P.J. (2008). Exploring network structure, dynamics, and function using NetworkX. In Proceedings of the 7th Python in Science Conference (SciPy 2008), pp. 11–15.

Haltiwanger, R.S., and Lowe, J.B. (2004). Role of Glycosylation in Development. Annu. Rev. Biochem. 73, 491–537.

Hamilton, W.L., Ying, R., and Leskovec, J. (2018). Inductive Representation Learning on Large Graphs. ArXiv:1706.02216 [Cs, Stat].

Henaff, M., Bruna, J., and LeCun, Y. (2015). Deep Convolutional Networks on Graph-Structured Data. ArXiv:1506.05163 [Cs].

Hu, W., Liu, B., Gomes, J., Zitnik, M., Liang, P., Pande, V., and Leskovec, J. (2020). Strategies for Pretraining Graph Neural Networks. ArXiv:1905.12265 [Cs, Stat].

Ichimiya, T., Nishihara, S., Takase-Yoden, S., Kida, H., and Aoki-Kinoshita, K. (2014). Frequent glycan structure mining of influenza virus data revealed a sulfated glycan motif that increased viral infection. Bioinformatics 30, 706–711.

Inoue, H. (2019). Multi-Sample Dropout for Accelerated Training and Better Generalization. ArXiv:1905.09788 [Cs, Stat].

Irie, A., Koyama, S., Kozutsumi, Y., Kawasaki, T., and Suzuki, A. (1998). The Molecular Basis for the Absence ofN-Glycolylneuraminic Acid in Humans. Journal of Biological Chemistry 273, 15866–15871.

Kapoor, A., Ben, X., Liu, L., Perozzi, B., Barnes, M., Blais, M., and O’Banion, S. (2020). Examining COVID-19 Forecasting using Spatio-Temporal Graph Neural Networks. ArXiv:2007.03113 [Cs].

Kightlinger, W., Warfel, K.F., DeLisa, M.P., and Jewett, M.C. (2020). Synthetic Glycobiology: Parts, Systems, and Applications. ACS Synth. Biol. 9, 1534–1562.

Koehler, M., Delguste, M., Sieben, C., Gillet, L., and Alsteens, D. (2020). Initial Step of Virus Entry: Virion Binding to Cell-Surface Glycans. Annu. Rev. Virol. 7, 143–165.

Lairson, L.L., Henrissat, B., Davies, G.J., and Withers, S.G. (2008). Glycosyltransferases: Structures, Functions, and Mechanisms. Annu. Rev. Biochem. 77, 521–555.

Lanteri, M., Giordanengo, V., Vidal, F., Gaudray, P., and Lefebvre, J.-C. (2002). A complete 1,3-galactosyltransferase gene is present in the human genome and partially transcribed. Glycobiology 12, 785–792.

Lauc, G., Kristic, J., and Zoldos, V. (2014). Glycans - the third revolution in evolution. Front. Genet. 5.

Letunic, I., and Bork, P. (2019). Interactive Tree Of Life (iTOL) v4: recent updates and new developments. Nucleic Acids Research 47, W256–W259.

Li, X., and Cheng, Y. (2020). Understanding the Message Passing in Graph Neural Networks via Power Iteration. ArXiv:2006.00144 [Cs, Stat].

Li, X., Xin, Y., Zhao, C., Yang, Y., and Chen, Y. (2020). Graph Convolutional Networks for Privacy Metrics in Online Social Networks. Applied Sciences 10, 1327.

Liu, K., Sun, X., Jia, L., Ma, J., Xing, H., Wu, J., Gao, H., Sun, Y., Boulnois, F., and Fan, J. (2019). Chemi-Net: A Molecular Graph Convolutional Network for Accurate Drug Property Prediction. IJMS 20, 3389.

van der Maaten, L., and Hinton, G. (2008). Visualizing data using t-SNE. Journal of Machine Learning Research 9, 2579–2605.

Manji, R.A., Lee, W., and Cooper, D.K.C. (2015). Xenograft bioprosthetic heart valves: Past, present and future. International Journal of Surgery 23, 280–284.

Merity, S. (2019). Single Headed Attention RNN: Stop Thinking With Your Head. ArXiv:1911.11423 [Cs].

Milewska, A., Zarebski, M., Nowak, P., Stozek, K., Potempa, J., and Pyrc, K. (2014). Human Coronavirus NL63 Utilizes Heparan Sulfate Proteoglycans for Attachment to Target Cells. Journal of Virology 88, 13221–13230.

Morris, C., Ritzert, M., Fey, M., Hamilton, W.L., Lenssen, J.E., Rattan, G., and Grohe, M. (2020). Weisfeiler and Leman Go Neural: Higher-order Graph Neural Networks. ArXiv:1810.02244 [Cs, Stat].

Nguyen, T., Nguyen, T., and Le, D.-H. (2020). Graph convolutional networks for drug response prediction. bioRxiv doi:10.1101/2020.04.07.030908.

Pang, L., Wang, M., Sun, X., Yuan, Y., Qing, Y., Xin, Y., Zhang, J., Li, D., and Duan, Z. (2018). Glycan binding patterns of human rotavirus P[10] VP8* protein. Virol J 15, 161.

Parker, R.B., and Kohler, J.J. (2010). Regulation of Intracellular Signaling by Extracellular Glycan Remodeling. ACS Chem. Biol. 5, 35–46.

Paszke, A., Gross, S., Massa, F., Lerer, A., Bradbury, J., Chanan, G., Killeen, T., Lin, Z., Gimelshein, N., Antiga, L., et al. (2019). PyTorch: An Imperative Style, High-Performance Deep Learning Library. ArXiv:1912.01703 [Cs, Stat].

Sarawagi, S., Chakrabarti, S., and Godbole, S. (2003). Cross-training: learning probabilistic mappings between topics. In Proceedings of the Ninth ACM SIGKDD International Conference on Knowledge Discovery and Data Mining - KDD ’03, (Washington, D.C.: ACM Press), p. 177.

Shelton, H., Ayora-Talavera, G., Ren, J., Loureiro, S., Pickles, R.J., Barclay, W.S., and Jones, I.M. (2011). Receptor Binding Profiles of Avian Influenza Virus Hemagglutinin Subtypes on Human Cells as a Predictor of Pandemic Potential. Journal of Virology 85, 1875–1880.

Solá, R.J., and Griebenow, K. (2009). Effects of glycosylation on the stability of protein pharmaceuticals. Journal of Pharmaceutical Sciences 98, 1223–1245.

Springer, S.A., and Gagneux, P. (2016). Glycomics: revealing the dynamic ecology and evolution of sugar molecules. Journal of Proteomics 135, 90–100.

Stanley, P. (2016). What Have We Learned from Glycosyltransferase Knockouts in Mice? Journal of Molecular Biology 428, 3166–3182.

Thompson, A.J., de Vries, R.P., and Paulson, J.C. (2019). Virus recognition of glycan receptors. Current Opinion in Virology 34, 117–129.

Torng, W., and Altman, R.B. (2019). Graph Convolutional Neural Networks for Predicting Drug-Target Interactions. J. Chem. Inf. Model. 59, 4131–4149.

Varki, A. (2017). Biological roles of glycans. Glycobiology 27, 3–49.

Viswanathan, K., Chandrasekaran, A., Srinivasan, A., Raman, R., Sasisekharan, V., and Sasisekharan, R. (2010). Glycans as receptors for influenza pathogenesis. Glycoconj J 27, 561–570.

Wu, F., Zhang, T., Souza Jr., A.H. de, Fifty, C., Yu, T., and Weinberger, K.Q. (2019). Simplifying Graph Convolutional Networks. ArXiv:1902.07153 [Cs, Stat].

Wu, Z., Pan, S., Chen, F., Long, G., Zhang, C., and Yu, P.S. (2020). A Comprehensive Survey on Graph Neural Networks. IEEE Trans. Neural Netw. Learning Syst. 1–21.

Yu, Y., Lasanajak, Y., Song, X., Hu, L., Ramani, S., Mickum, M.L., Ashline, D.J., Prasad, B.V.V., Estes, M.K., Reinhold, V.N., et al. (2014). Human Milk Contains Novel Glycans That Are Potential Decoy Receptors for Neonatal Rotaviruses. Molecular & Cellular Proteomics 13, 2944–2960.

Zhao, Y.-Y., Takahashi, M., Gu, J.-G., Miyoshi, E., Matsumoto, A., Kitazume, S., and Taniguchi, N. (2008). Functional roles of N-glycans in cell signaling and cell adhesion in cancer. Cancer Science 99, 1304–1310.

