## Supplemental Figures for "Using Graph Convolutional Neural Networks to Learn a Representation for Glycans"

### Supplementary Figures

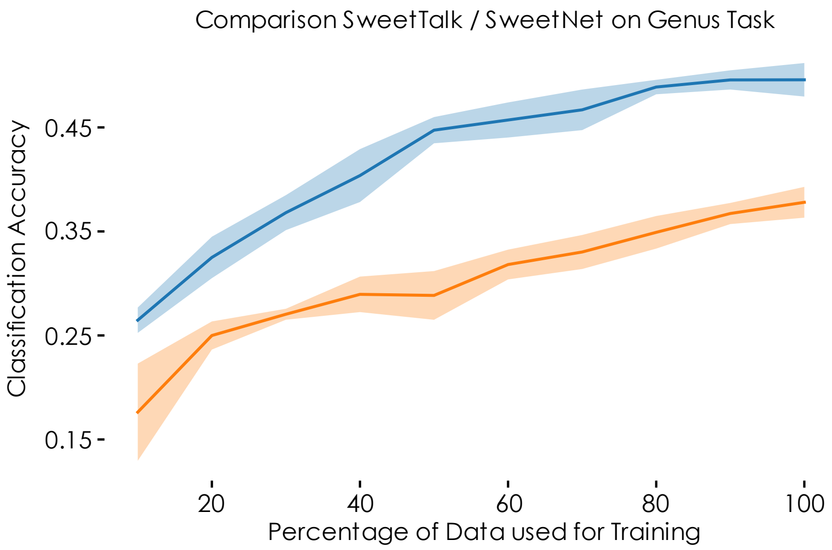


**Supplementary Figure 1. Model comparison regarding data efficiency; Related to Table 1.** SweetNet-based (blue) and SweetTalk-based (orange) models were trained to predict the genus a given glycan sequence stemmed from. We trained these models with a random selection of 10-100% of the training dataset and evaluated the resulting model on a separate validation set comprising 20% of the overall data prior to selecting the training data. The data shown are the means (line) and standard deviations (shaded area) of five training runs for each percentage of the training data we tested.

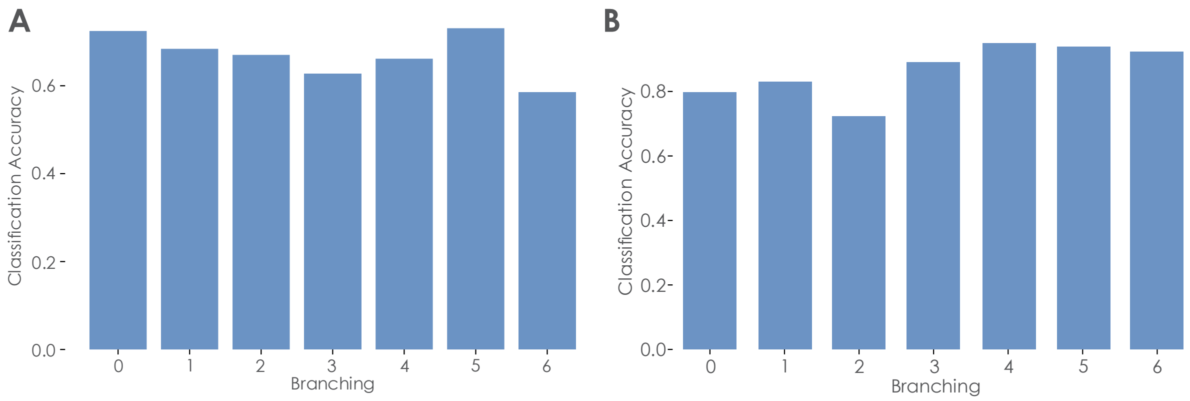


**Supplementary Figure 2. The influence of branching on model performance; Related to Table 1. (A-B)** Assessing model performance at every level of branching. For the trained SweetTalk-based (**A**) and SweetNet-based (**B**) models to predict the taxonomic class of a glycan, we depict the averaged prediction accuracy on the entire dataset for every level of glycan branching up to a branching of six, as higher levels of branching were too rare in our dataset for firm conclusions.

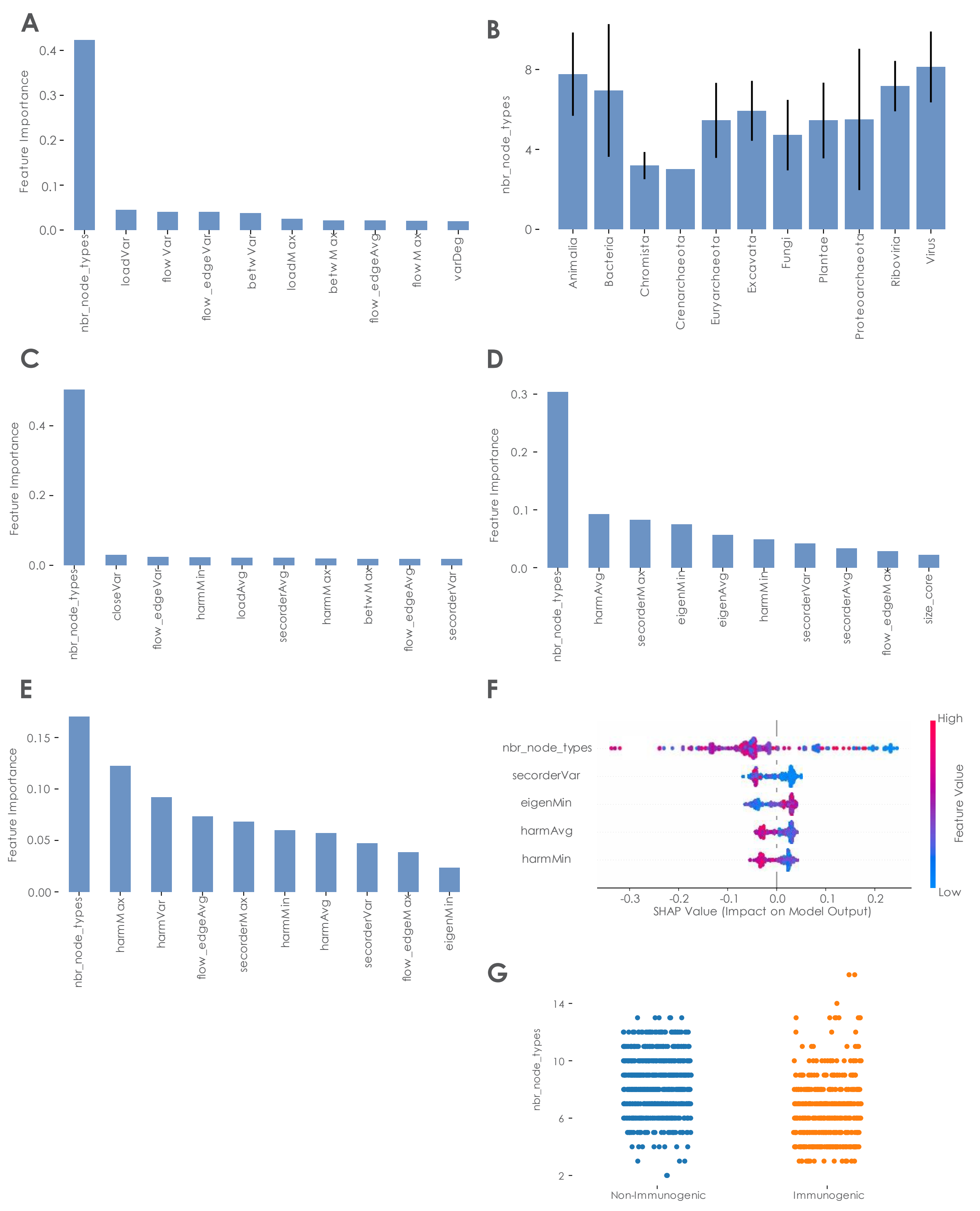


**Supplementary Figure 3. Graph properties important for glycan classification; Related to Figure 2. (A)** Feature importance for a Random Forest model trained to predict the taxonomic kingdom of a glycan. The 10 most important features are shown. (**B**) Distribution of nbr_node_types across taxonomic kingdoms. Means and standard deviations of nbr_node_types for each taxonomic kingdom are shown. (**C-E**) Feature importance for a Random Forest model trained to predict the contribution of a glycan to pathogenicity (**C**), glycan immunogenicity (**D**), and glycan class (**E**). The 10 most important features are shown. (**F**) SHAP (SHapley Additive exPlanations) values for important variables of the glycan immunogenicity prediction model. The effect of variable values on the prediction is shown. (**G**)

Distribution of number of node types between non-immunogenic and immunogenic glycans. Using a Welch's t-test, we ascertained that the difference is highly significant (p-value: 1.47e-45).

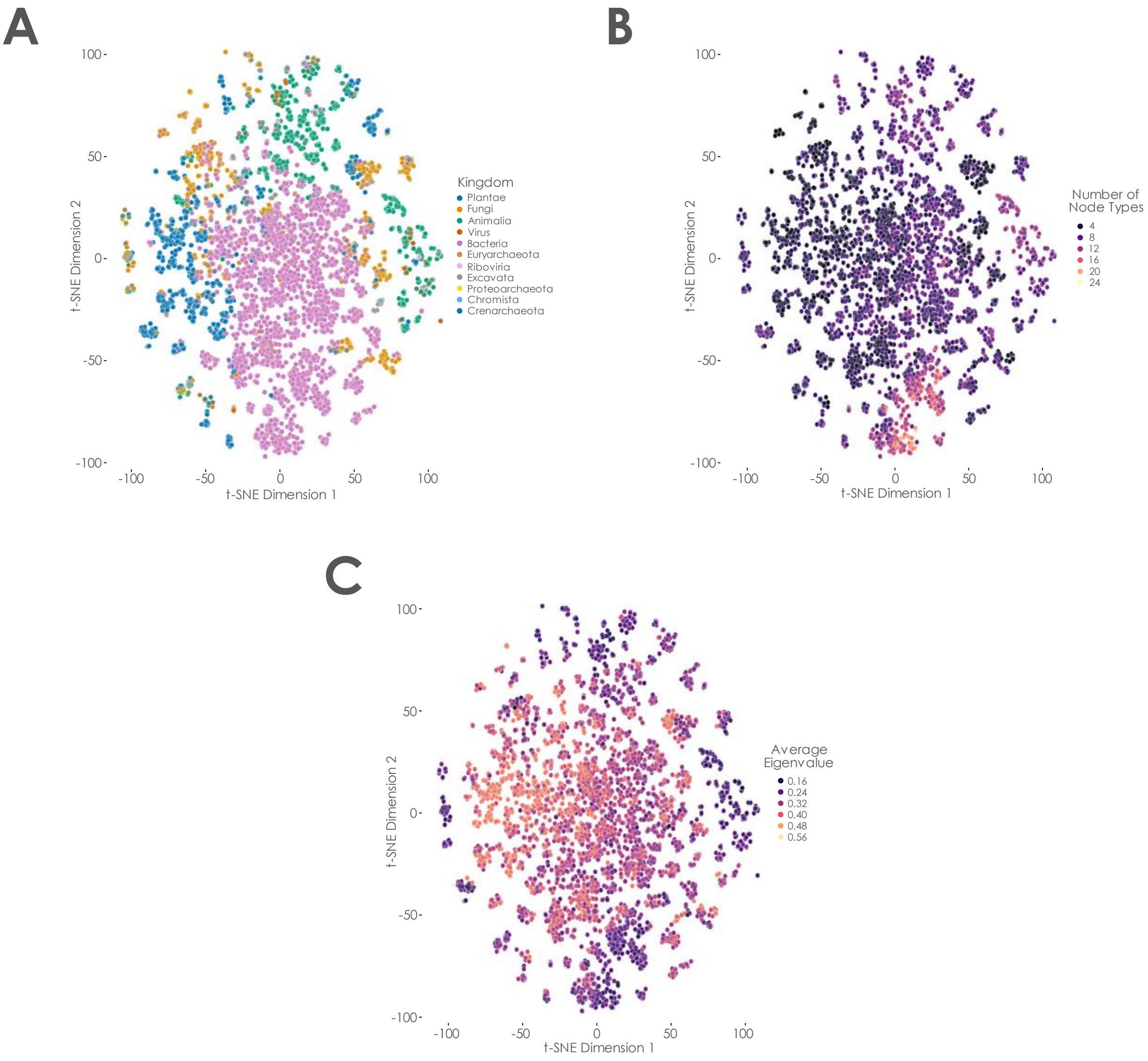


**Supplementary Figure 4. Clusters of glycans with certain graph properties in SweetNet representations; Related to Figure 2. (A-C)** Glycan representations for all glycans with taxonomic information in our dataset were generated by using them as input for our trained kingdom-level SweetNet model and extracting the representation learned by the model, shown here via t-SNE. We then colored glycans according to their taxonomic kingdom (**A**), number of node types (**B**), or average eigenvalues (**C**).
